# The spectral underpinnings of pathogen spread on animal networks

**DOI:** 10.1101/2022.07.28.501936

**Authors:** Nicholas M. Fountain-Jones, Mathew Silk, Raima Carol Appaw, Rodrigo Hamede, Julie Rushmore, Kimberly VanderWaal, Meggan E Craft, Scott Carver, Michael Charleston

## Abstract

Predicting what factors promote or protect populations from infectious disease is a fundamental epidemiological challenge. Social networks, where nodes represent hosts and edges represent direct or indirect contacts between them, are important in quantifying these aspects of infectious disease dynamics. However, how network structure and epidemic parameters interact in empirical networks to promote or protect animal populations from infectious disease remains a challenge. Here we draw on advances in spectral graph theory and machine learning to build predictive models of pathogen spread on a large collection of empirical networks from across the animal kingdom. We show that the spectral features of an animal network are powerful predictors of pathogen spread for a variety of hosts and pathogens and can be a valuable proxy for the vulnerability of animal networks to pathogen spread. We validate our findings using interpretable machine learning techniques and provide a flexible web application for animal health practitioners to assess the vulnerability of a particular network to pathogen spread.

## Introduction

Capturing patterns of direct or indirect contacts between hosts is crucial to model pathogen spread in populations (Newman 2002; Craft 2015; Pastor-Satorras *et al*. 2015; Sah *et al*. 2018, 2021). Increasingly, contact or social network approaches, where hosts are nodes and edges reflect interactions between hosts, play a central role in epidemiology and disease ecology (e.g., Meyers *et al*. 2005; Bansal *et al*. 2007; Eames *et al*. 2015; White *et al*. 2017). When there is heterogeneity in contacts between individuals, network approaches can provide more nuanced and reliable estimates of pathogen spread, including in wildlife populations (e.g., Meyers *et al*. 2006; Bansal *et al*. 2010; Craft *et al*. 2011). Formulating general rules for how easy-to-calculate network structure properties may promote or protect populations from pathogen spread can reveal important insights into how host behaviour can mediate epidemic outcomes (Sah *et al*. 2017), and provide practitioners with a proxy for how vulnerable a population is to disease without extensive simulations (Silk *et al*. 2017; Sah *et al*. 2018). Further, structural properties of networks can be incorporated into traditional susceptible– infected–recovered (SIR) models to account for contact heterogeneity when predicting pathogen dynamics across populations (e.g., Meyers *et al*. 2005; Bansal *et al*. 2007; Pastor-Satorras *et al*. 2015). For example, approaches such as degree-based mean field approximation (DBMF), can estimate outbreak size using the degree distribution of uncorrelated networks (i.e., the degree of one node is uncorrelated with neighbouring nodes, (Newman 2002; Pastor-Satorras *et al*. 2015)).

However, it remains unclear whether one structural characteristic or a combination of network characteristics can reliably predict pathogen dynamics across empirical networks in different biological systems (Ames *et al*. 2011; Sah *et al*. 2018). For example, species that are more social tend to have more clustered or “modular” networks, and this modularity has been found to increase (Lentz *et al*. 2012), reduce (Salathé & Jones 2010), or have little effect (Sah *et al*. 2018) on outbreak size across different systems. The average number of contacts between hosts can be identical across networks and yet still result in substantially different outbreak patterns (Ames *et al*. 2011). Even the apparent size (number of nodes) of the network, often constrained by limitations of sampling, can impact estimates of pathogen spread, particularly in wildlife populations (McCabe & Nunn 2018). As network characteristics such as network size and modularity are often correlated (Newman 2006; Silk *et al*. 2017) and can have complex impacts on spread (Sah *et al*. 2017; McCabe & Nunn 2018; Porter 2020), determining those characteristics that affect outbreaks remains a fundamental question in infectious disease biology (Sah *et al*. 2018).

Searching for general relationships between network structure and pathogen spread in animal populations is further challenged, as the relationship is also affected by host-pathogen traits, such as infectiousness and recovery rate. For example, modularity appears to make no difference to pathogen prevalence for highly infectious pathogens (Sah *et al*. 2017). Pathogens with long recovery rates can increase outbreak size across networks (Shu *et al*. 2016). Given that we rarely have reliable estimates of pathogen traits in wild populations (e.g., for different probabilities of infection per contact, or recovery rates), any predictive model of the relationship between spread and network structure would ideally be generalizable across pathogens.

Advances in spectral graph theory have the potential to resolve this challenge, offering an additional set of measures based on the **spectrum** of a network rather than average node or edge level attributes (see Text Box 1 for further definitions of terminology in bold). A graph spectrum is the set of **eigenvalues** (often denoted with a Greek lambda λ) of a matrix representation of a network. Theoretical studies have shown relationships between particular eigenvalues and connectivity across networks are independent of pathogen propagation models (Prakash *et al*. 2010). For example, networks with a high algebraic connectivity, also known as the **Fiedler value** (the second smallest eigenvalue of the network’s **Laplacian matrix**) are “more connected” than those with low values. It has been found that, in ecological networks for example, if the Fiedler value is sufficiently large, removing edges will have little effect on overall network connectivity (Kumar *et al*. 2019), but whether this lack of effect is mirrored by pathogen dynamics is not yet clear. Another quantity of interest is the **spectral radius** – the largest absolute value of the eigenvalues of its **adjacency matrix**. The link between the spectral radius and epidemiological dynamics is better understood, with theoretical work showing that this value closely mirrors both epidemic behaviour and network connectivity (Prakash *et al*. 2010) and has been used to understand vulnerability of cattle networks to disease (Darbon *et al*. 2018). For example, networks with the same number of edges and nodes but higher spectral radius (λ_1_) are more vulnerable to outbreaks than networks with low spectral radius (λ_1_→1). We hypothesize that spectral measures such as these have great potential to improve our ability to predict dynamics of pathogen spread on networks, where previous methods such as modularity have proved inadequate (Sah *et al*. 2017).

#### Text Box 1: Terminology used in this paper.

A *graph* (or “*network*”) is a collection of *nodes* and a collection of *edges* connecting the nodes in pairs, e.g., nodes *x, y* joined by edge (*x*,*y*). We define the *size* of the network – usually *n*, as the number of nodes (this usage differs from other strict mathematical definitions, but we feel this is more intuitive). Two nodes are said to be *adjacent* if they are connected by an edge, and the number of vertices adjacent to a given vertex *x* is called its *degree*, deg(*x*). Edges may be directed, in which case edge (*x*,*y*) is different from edge (*y*,*x*), but in our analyses we treat them as *undirected*, so (*x*,*y*)=(*y*,*x*). Graphs can be represented naturally by matrices whose rows and columns are indexed by the nodes (1, 2,…, *n*): the obvious one is the *adjacency matrix A*, whose (*i*,*j*)-th entry *A_ij_* is 1 if nodes *i* and *j* are adjacent, and 0 otherwise. *A* is symmetric and *n* × *n*, as are all the matrices in this work. Another useful matrix is the *degree* matrix *D*, in which *D_ij_* is the degree of node *i* if *i*=*j*, and 0 otherwise. The *Laplacian* matrix *L* is the most complex one we use herein, but is easily calculated using *L_ij_* = *D_ij_* – *A_ij_*.

The *eigenvalues* of a matrix are solutions to the matrix equation *M**v*** = λ***v***, where *M* is a matrix and ***v*** a vector of the appropriate size. Solving for ***v*** yields λ. These eigenvalues, ordered by their size, form the *spectrum* of a graph, as derived using any of the matrices just described. The *Fiedler value* of a graph is the second-smallest eigenvalue of *L*, and the s*pectral radius* is the largest eigenvalue of *A*.

Measures of *Modularity* such as the Newman *Q* coefficient capture the strength of division within a network by quantifying the density of edges within and between subgroups. When there is no division within the network as the density of edges is the same between and within subgroups *Q* = 0, whereas higher values of *Q* indicate stronger divisions (Newman 2006). As *Q* scales with network size (small networks being generally less modular), relative modularity (*Q_rel_*) allows for comparison across network sizes by normalizing Q using the maximum possible modularity for the network (*Q_max_*) (Sah et al. 2017).

We assess the predictive capability of spectral values compared to other structural attributes such as **modularity** (Newman’s Q; (Newman 2006)) using advances in machine learning to construct non-linear models of simulated pathogen spread across a large collection of empirical animal networks, including those from the Animal Social Network Repository (ASNR) (Sah *et al*. 2019). The ASNR is a large repository of empirical social networks that provides novel opportunities to test the utility of spectral values in predicting spread across a wide variety of, mostly animal, taxa across a diversity of social systems -- from eusocial ants (Arthropoda: Formicidae) to more solitary species such as the desert tortoise (*Gopherus agassizii*). We combined networks from this resource with other published networks, including badgers (*Meles meles*) (Weber *et al*. 2013), giraffes (*Giraffa camelopardalis*) (VanderWaal *et al*. 2014), and chimpanzees (*Pan troglodytes*) (Rushmore *et al*. 2013) to generate a dataset of over 600 unweighted networks from 51 species. We treated each network as static (i.e., edges are fixed for the duration of the simulated outbreak). While dynamic networks where contact patterns fluctuate over time can provide more realistic estimates of pathogen spread (Bansal *et al*. 2010), they are rare, particularly for wild populations. Therefore, we were restricted to analysing static networks, though further study of dynamic networks would be valuable. We then simulated pathogen spread using a variety of SIR parameters and harnessed recent advances in multivariate interpretable machine learning models (*MrIML;* (Fountain-Jones *et al*. 2021)) to construct predictive models. As many species were represented by multiple networks, often over different populations and or timepoints and constructed in different ways (e.g., some edges reflected spatial proximity rather than direct contact), we included species and network construction variables in our models to account for these correlations in addition to exploring the diversity of network structures across the animal kingdom.

We demonstrate that using spectral measures of network structure alone can provide a useful proxy for network vulnerability. Our interpretable machine learning models identify putative threshold values for the vulnerability of a network to pathogen spread. Further, we provide a user-friendly application that utilizes our models to provide practitioners with predictions, for example, of the prevalence of a pathogen across a variety of spread scenarios using a user-supplied network without the need for lengthy simulation (see https://spreadpredictr.shinyapps.io/spreadpredictr/).

## Methods

### Networks

We downloaded all animal social networks from the ASNR on 12^th^ January 2022 (Sah *et al*. 2019) and combined these with other comparable published animal networks (Rushmore *et al*. 2013; Weber *et al*. 2013; VanderWaal *et al*. 2014). These networks included those reflecting direct and indirect (such as burrow sharing) contacts between hosts (hereafter ‘contact networks’). Farmed domestic animals were not included in our analyses. We binarized each network, extracted the largest connected component, and excluded networks with fewer than 10 individuals. This left us with 603 networks from 43 species.

From each network we calculated a variety of network structure variables using the R package *igraph* (Csárdi & Nepusz 2006) (see Table S1 for a summary of the variables measured). As these networks were constructed using different techniques, we also extracted metadata from the ASNR or the publication associated with the network (Table S2). These variables were also added to the models. We used Principal Components Analysis (PCA) biplots to examine the drivers of variation in network structure and visualise how networks clustered by taxonomic class. We removed eight networks with missing metadata and screened for correlations between variables. We used a pairwise correlation threshold of 0.7 to remove the variable from each such pair with the highest overall correlation. We used this relatively high threshold as many of the machine learning algorithms are less sensitive to variable collinearity (Fountain-Jones *et al*. 2019)

### Simulations

To simulate the spread of infection on each network we used our R package “EpicR” (<u>Epi</u>demics by <u>c</u>omputers in <u>R</u>; available on GitHub at https://github.com/mcharleston/epicr). The simulations use a standard discretisation of the SIR model, in which time proceeds in “ticks,” for example representing days. Initially one individual was chosen at uniform random to be infected (I) and all others were susceptible (S). At each time step, one of two changes of state can happen to each individual (represented by a node), depending on its current state. An ‘S’ individual will become infected (I) with a probability (1 − (1 − β)*^k^*), where *k* is the number of currently infected neighbours it has, or otherwise stay as S; an ‘I’ individual will recover (R) with probability γ or remain as I. Recovered (R) individuals stay as R.

In classic deterministic SIR models as a set of differential equations, β and γ are instantaneous rates; here, they are probabilities per time step, so at a coarse level, they are comparable.

On each network, we performed 1000 simulations using different combinations of transmission (β = 0.01, 0.025, 0.05, 0.1, 0.2) and recovery probabilities (γ = 0.02, 0.04, 0.2, 0.4). We chose these SIR parameter values to broadly reflect a range of scenarios from high to low transmissibility and slow to fast recovery (Leung 2021) and ensure large outbreaks (>10% of individuals infected, (Sah *et al*. 2017)). One randomly chosen individual was infected at the beginning of the simulation and we ran the simulation for 100 time steps to ensure that the epidemic ended and there were no remaining infected nodes. For each simulation we recorded two complementary epidemic measures to capture disease burden and speed of spread: a) the maximum prevalence reached, or the maximum proportion of individuals infected in the network after 100 time steps (hereafter ‘proportion infected’) and b) time to outbreak peak (i.e., which time step had the maximum number of infections, hereafter ‘time to peak’). The epidemic measures estimated in the simulations were used as the response variables in the machine learning models.

### Machine learning pipeline

We used a recently developed multi-response interpretable machine learning approach (*MrIML*) (Fountain-Jones *et al*. 2021) to predict outbreak characteristics using network structure variables. Our *MrIML* approach had the advantage of allowing us to rapidly construct and compare models across a variety of machine-learning algorithms for each of our response variables as well as assess generalized predictive surfaces across epidemic parameters.

To test the robustness of our results, we compared the performance of four different underlying supervised regression algorithms in our *MrIML* models. We compared linear models (LMs), support vector machines (SVMs), random forests (RF) and gradient boosted models (GBMs) as they operate in markedly different ways that can affect predictive performance (Fountain-Jones *et al*. 2019; Machado *et al*. 2019). For the best performing model, we also tested algorithm performance using just the spectral radius and Fiedler value for each network. Categorical predictors such as ‘species’ were hot-encoded for some models as needed (i.e., representing the variable as a binary vector, see Table S4). As proportion infected and time to peak are continuous response variables, we compared the performance of each algorithm using the average R^2^ and root mean squared error (RMSE) across all responses (hereafter, the ‘global model’). As we included models that were not fitted using sums of squares, our R^2^ estimate depended on the squared correlation between the observed and predicted values (Kvålseth 1985). As ants (Insecta: Formicinae) were over-represented, we compared model performance and interpretation with and without these networks using 10-fold cross validation, to prevent overfitting. We tuned hyperparameters for each model (where appropriate) using 100 different hyper-parameter combinations (a 10×10 grid search) and selected the combination with the lowest RMSE. The underlying algorithm with the highest predictive performance was interrogated further.

We interpreted this final model using a variety of model-agnostic techniques within the *MrIML* framework. We assessed overall and model-specific variable importance using a variance-based method (Greenwell *et al*. 2018). We quantified how each variable alters epidemic outcomes using accumulated local effects (ALEs) (Apley & Zhu 2016). In brief, ALEs isolate the effect of each network characteristic on epidemic outcomes using a sliding window approach, calculating the average change in prediction across the values’ range (while holding all other variables constant) (Molnar 2018). ALEs are less sensitive to correlations and straightforward to interpret as points on the ALE curve are the difference in the prediction of each observation from the mean prediction (Apley & Zhu 2016; Molnar 2018; Fountain-Jones *et al*. 2021).

To further examine the predictive performance of our black-box models (SVM, RF and GBM) we calculated a global surrogate decision tree (hereafter ‘global surrogate’) to approximate the predictions of our more complex trained models. Global surrogates are generated by training a simpler decision tree to the *predictions* (instead of observations) of the more complex ‘black box’ models using the network structure data. How well the surrogate model performed compared to the complex model is then estimated using R^2^. See Molnar (2018) for details.

We gained more insight into model behaviour and how network structure impacted epidemic outcomes on individual networks, including by calculating Shapley values (Štrumbelj & Kononenko 2014). Shapley values use a game theoretic approach to play off variables in the model with each other based on their contribution to the prediction (Shapley 1953). Calculating Shapley values involves solving a system of linear equations to assign each feature a unique weight based on its contribution to the predicted output and its interaction with other features (Štrumbelj & Kononenko 2014). For example, negative Shapley values indicate that the observed value ‘contributed to the prediction’ by reducing the proportion infected or time to peak in an outbreak for a particular network. See Molnar (2018) for a more detailed description and Fountain-Jones *et al*. (2019) and Worsley-Tonks *et al*. (2020) for how Shapley values can be interpreted in epidemiological settings. See https://github.com/nfj1380/igraphEpi for our complete analytical pipeline.

Lastly, we also compared our machine learning estimates of proportion infected to those derived from the DBMF SIR approximations for each network. Briefly, the DBMF approximation calculates the proportion infected using the following equation:

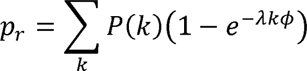

Where P(k) is the degree distribution of the network, λ is the epidemic threshold and k is the first moment of the degree distribution. The auxiliary function (Φ) cannot be solved analytically, but can be approximated using the network epidemic transition threshold (λ_c_):

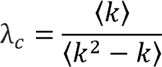

where k^2^ is the second moment of the degree distribution. See Pastor-Satorras *et al*. (2015) for more details.

## Results

### Diversity of network structures

We identified substantial variation in network structure across animal taxa. The static unweighted animal social networks in our database ranged from nearly completely unconnected (Spectral radius λ_1_ ∼ 1, Fiedler value ∼0, not included in our predictive models) to highly connected (Spectral radius λ_1_ ∼ 160, Fiedler value ∼ 140, Fig. 1). Similarly, the networks ranged from homogeneous (i.e., not modular, Q_rel_ = 0, see Text Box 1) to highly modular and subdivided (Q_rel_ > 0.8). Our principal component analysis (PCA) identified key axes of structural variation across empirical networks (Fig. 2). The first principal component (PC1) distinguished networks that had a large diameter and mean path length and were highly modular (PC1 < 0) from networks with a high mean degree and transitivity (PC1 > 0, Fig. 2, see Table S3). The second principal component (PC2) separated networks based on network size (number of nodes), maximum degree and the network duration (i.e., the time period over which the network data was collected, Fig. 2). The eusocial ant networks (*Camponotus fellah*, Insecta: Hymenoptera) and mammal networks tended to cluster separately (Fig. 2), with the other taxonomic classes dispersed between these groups (Fig. 2) or species (see Fig. S1 for clustering by species). The networks’ spectral properties (the Fiedler value and spectral radius) explained a unique portion of structural variance that did not covary with other variables (see Table S3 for vector loadings and Fig S2 for all pair-wise correlations). We found variables such as mean degree and transitivity the most correlated with the other variables and were excluded from further analysis (Tables S2, Fig S2).

**Fig. 1:**
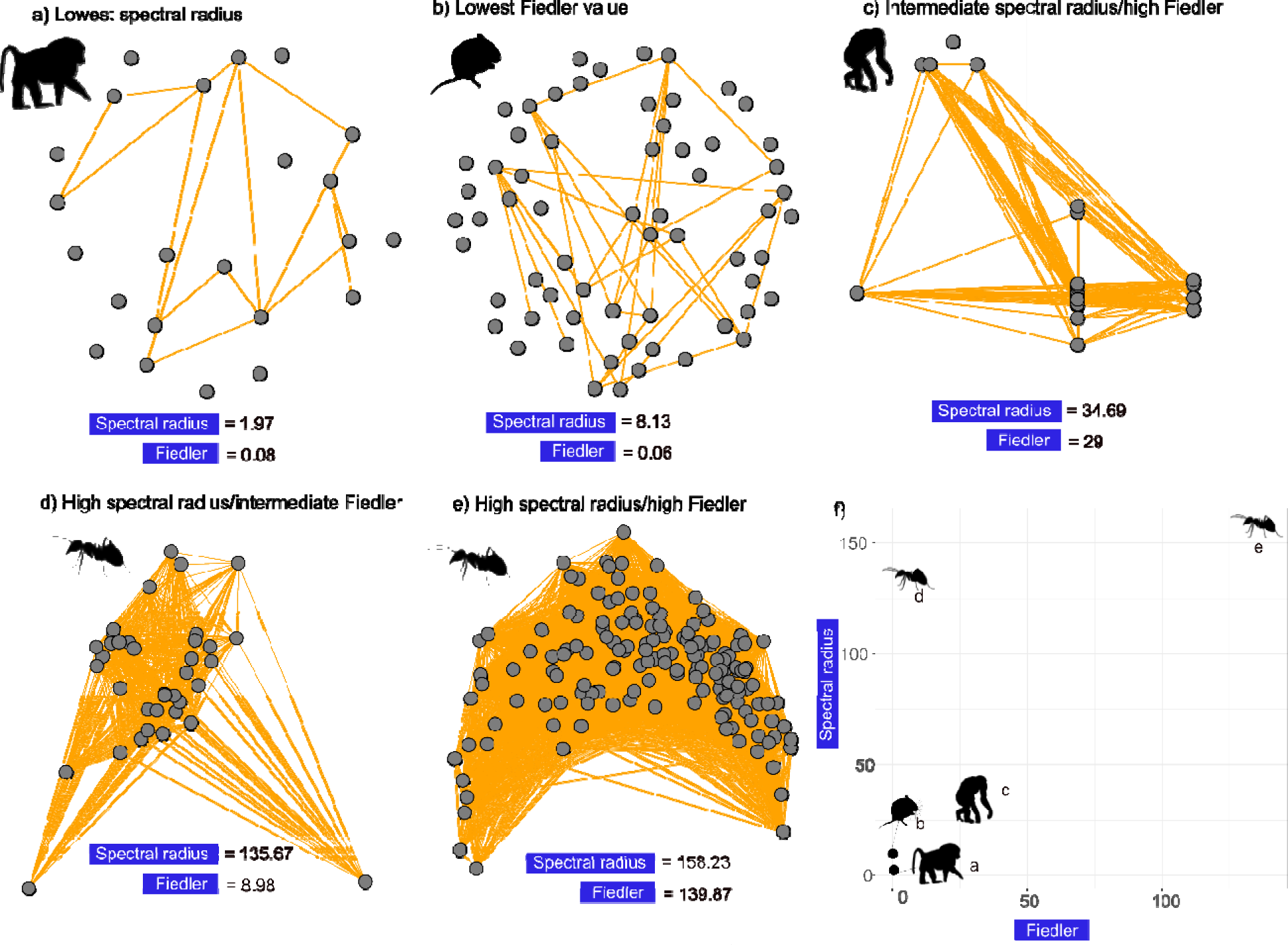
Examples of networks analysed in this study with a) the lowest spectral radius (baboon *Papio cynocephalus* contact network), b) the lowest Fiedler value (vole *Microtus agrestis* trap sharing network), c) intermediate spectral radius values but high Fiedler value (Chimpanzee *Pan troglodytes* contact network), d) high spectral radius/intermediate Fiedler value (*Camponotus fellah* colony contact network) and e) high values of both measures (another *C. fellah* colony contact network). The mean values across all networks were 34.80 and 7.31 for the spectral radius and Fiedler value respectively. f) summary of values across networks (a-e). Silhouettes were sourced from phylopic (http://phylopic.org/). Note that disconnected nodes were not included in the analysis.

**Fig. 2:**
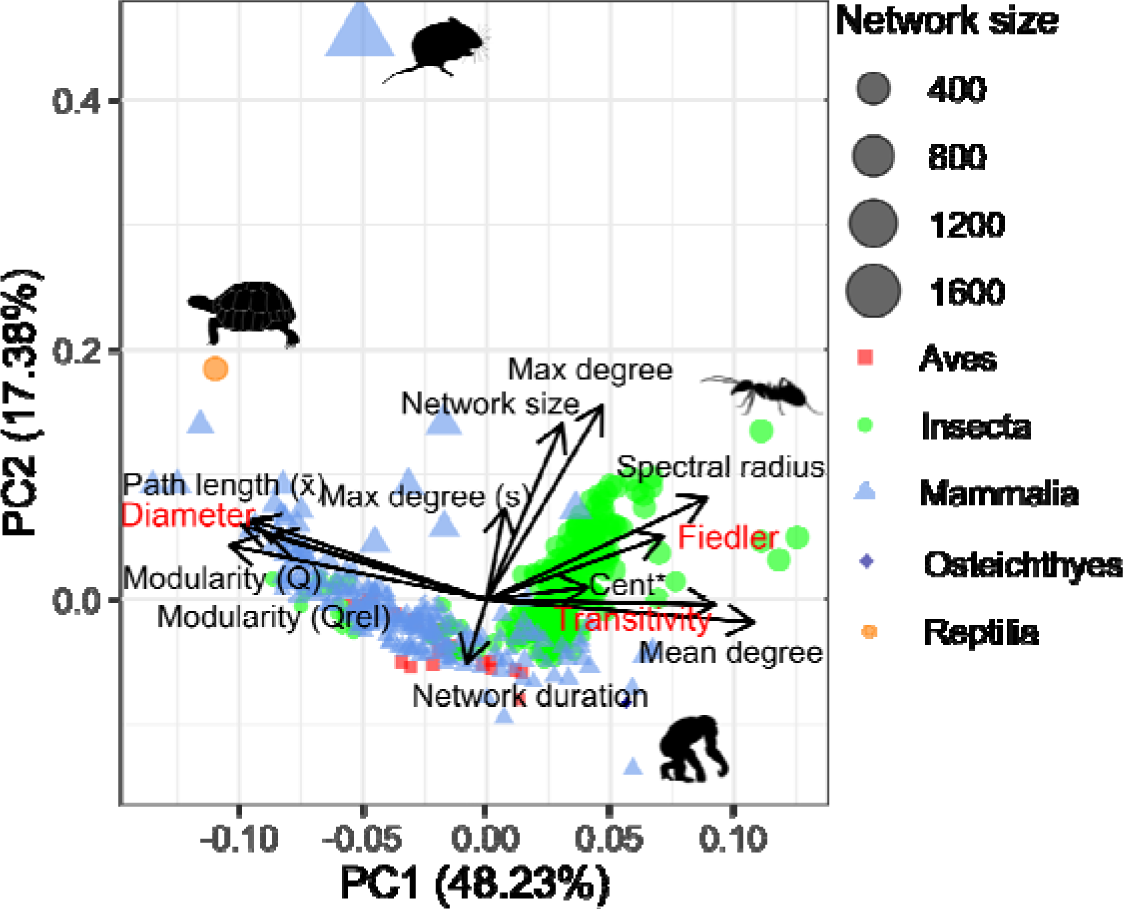
Principal components analysis (PCA) biplot showing that network structure largely clusters by taxonomic class. Points are coloured by taxa. Points closer together in Euclidean space have networks more similar in structure. Points are scaled by network size. The length and direction of vectors (black arrows) shows how each variable relates to each principal component with larger vectors having higher loadings on that axis. The PCA was constructed just using continuous network characteristics. Percentages next to PC scores indicate how much variability in the data is accounted for by each axis. See Table S1 for axis loadings and Fig. S1 for the species-level clustering. See Tables S2 & S3 for variable definitions. Silhouettes for some of the outlying networks were sourced from PhyloPic (http://phylopic.org/). s = scaled. Cent = Centralization.

### Spectral properties predict pathogen spread across epidemic scenarios

Across all SIR parameter combinations, spectral measures played a dominant role in our predictive models (Fig. 3a, Fig. S3). We could predict the proportion infected in a network and time to peak using both spectral measures and species identity alone (Table S4). To aid interpretation we present the analyses from two recovery probabilities (γ = 0.04, 0.4 see Fig S4 for the analysis with a wider variety of recovery probabilities). Network size, relative modularity and centralization, for example, were less important in predicting proportion infected across all SIR model parameter combinations tested (Fig. 3a). Nonlinear relationships were likely important for prediction of proportion infected, as random forests (RF) had the highest predictive performance overall (Table S4) and substantially outperformed linear regression in the *MrIML* framework (root mean square error (RMSE) 0.03 vs 0.13). Our RF model just including the spectral radius and Fiedler value had very similar predictive performance (R^2^, RMSE) compared to the models with all variables included (Table S4). Our models also substantially outperformed DBMF approximations with a ∼ten-fold decrease in RMSE across a range of β and γ values (Table S5).

**Fig. 3:**
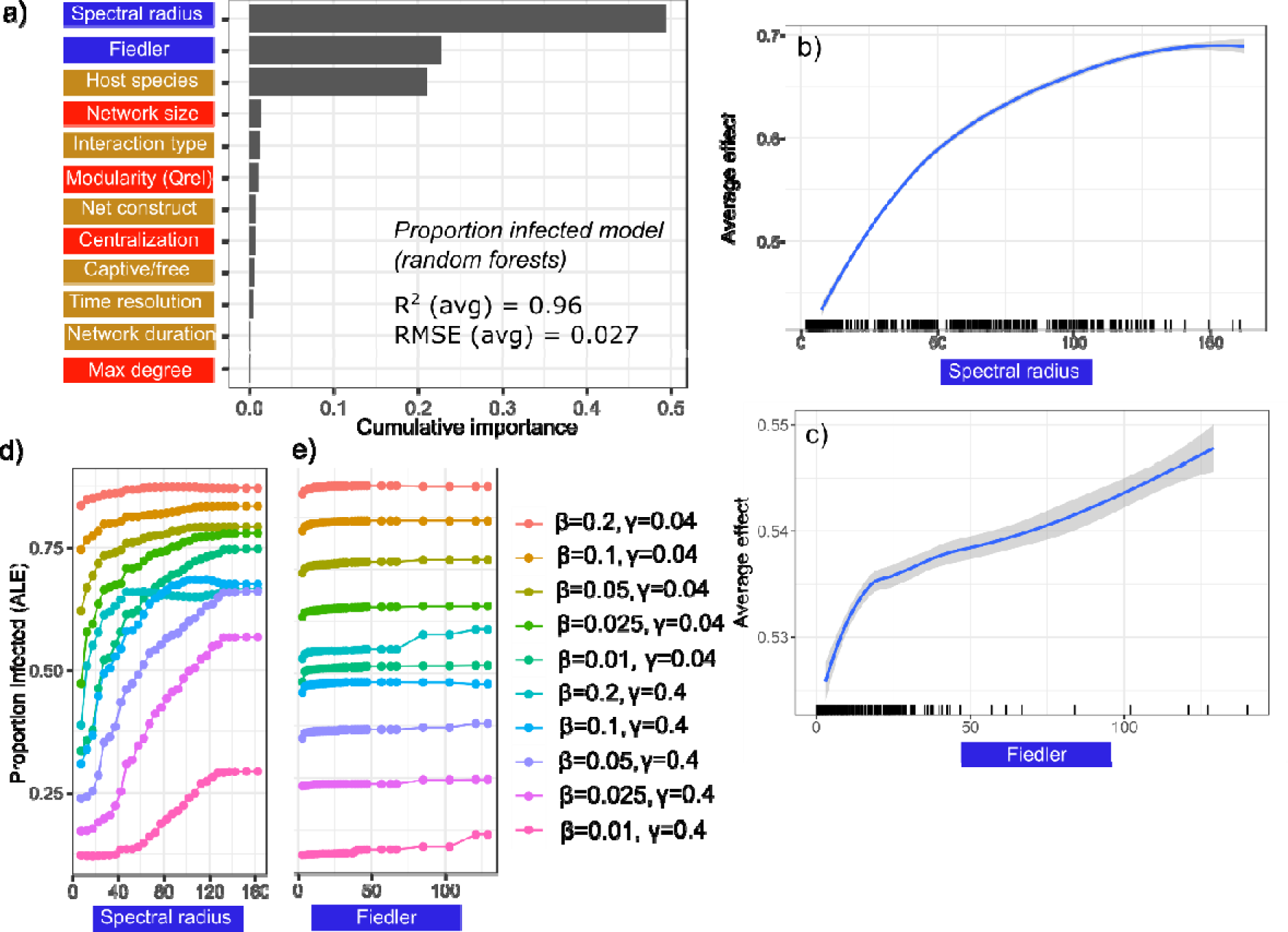
Plots showing the predictive performance, variable importance and the functional form of relationships for our best-performing *MrIML* proportion infected model. See Table S4 for model performance estimates across algorithms. The colour of the labels indicates what type of predictor it is (blue = spectral, red = non-spectral network structural variables, gold = network metadata, see Tables S1 & S2). a) Spectral radius and the Fiedler value (followed by species) are the most important predictors of proportion of individuals infected across all simulations (importance threshold >0.1) and overall model performance was high (average R^2^ = 0.96 and root mean square error (RMSE) = 0.027). b-c) Average predictive surface showing the relationship between spectral properties and proportion infected across all epidemic values (95% confidence intervals in grey). Rug plot on the x axis of the panels on the right shows the distribution of each characteristic across empirical networks. d-e) The accumulated local effects (ALE) plot revealed that the strongly non-linear relationships between both spectral properties and proportion infected were mediated by transmission and recovery probabilities. We chose these SIR parameter values (β = transmission probability, γ = recovery probability) to ensure major outbreaks occurred on the empirical networks. Net construct = Network construction method.

Variable importance and predictor conditional effects were consistent between the machine learning algorithms, so we subsequently analysed the best performing RF model. We found a nonlinear relationship between proportion infected and spectral radius across each SIR parameter combination, with the average prediction of proportion infected increasing by ∼30% across the range of spectral radius values (holding all other variables constant in the model, Fig. 3b). In contrast we found a more modest effect of the Fiedler value, with the proportion of infected only increasing on average ∼3% across the observed range of values for all SIR parameters (β & γ, Fig 3c). When the Fiedler value was greater than about 15, we did detect a minor increase in the proportion infected in networks (Fig. 3c). However, there was variation in the relationship between the proportion infected and these spectral values across transmission (β) and recovery probabilities (γ, Figs. 3d-e). For example, when the probability of transmission was relatively high (β = 0.2) and recovery low (γ = 0.04) the proportion infected across networks was ∼80% and spectral radius had a relatively minor effect (Fig. 3d). A network’s spectral radius had a stronger effect when the probability of recovery was higher (γ = 0.4) across all values of β. The increase in proportion infected when the Fiedler value was low (< 15) was not apparent when spread was slower and chances of recovery higher (e.g., β = 0.025 or 0.01, γ = 0.4; Fig 3e). The spectral radius and Fiedler value patterns overall were similar, with larger values reducing the time-to-peak (Fig. S3). However, modularity played a greater role in our time to peak models, with the time to peak being longer for more modular networks above a Q_rel_ threshold of ∼ 0.75 (Fig. S5).

### Simplifying our models with global surrogates

When we further interrogated our moderate (β = 0.05) transmission models, we found that the spectral radius and Fiedler value overall also played a dominant role in our predictions of spread. To quantify the putative mechanisms that underlie our model predictions – ‘to decloak the black box’ − and gain insight into possible interactions between predictors, we constructed surrogate decision trees as a proxy for our more complex RF model. We trained our surrogate decision tree on the predictions of the RF model rather than the network observations directly. In each case, the surrogate decision tree approximated the predictions of our models (thousands of decision trees) remarkably well (global R^2^ > 0.95, see (Molnar 2018) for details). The spectral radius and, to a lesser extent, the Fiedler value and modularity values dominated surrogate trees for all SIR parameter sets (Fig. 5, Figs. S6 & S7). For example, for networks with a Fiedler value ≥ 0.86 and a spectral radius ≥ 20 (as was the case for 51% of our networks, Fig. 4b) the estimated maximum proportion of the network infected was 0.92 (Fig. 4b). The duration over which the data was collected also was included in the surrogate model, with networks collected over > 6.5 days having higher estimates of proportion infected (Fig. 4).

**Fig. 4.**
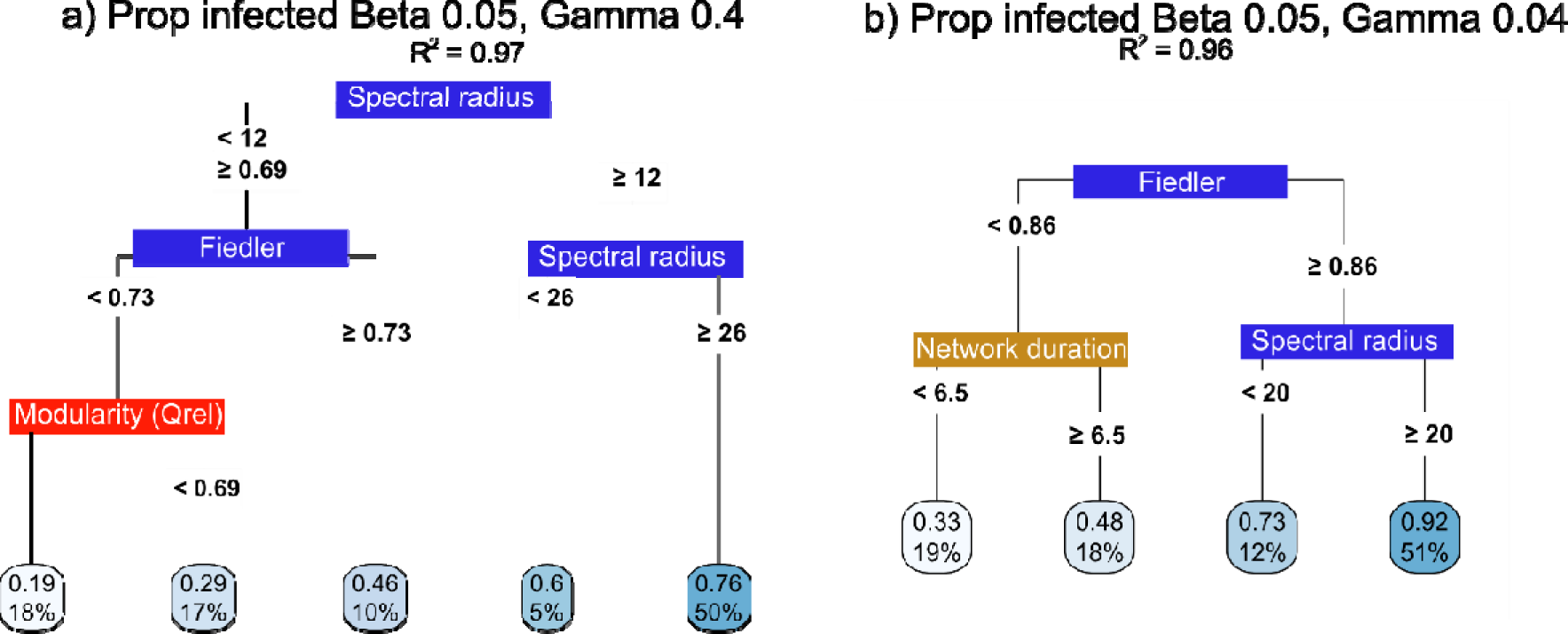
Global surrogate decision trees for our moderate transmission (β = 0.05) proportion infected with a) high and b) low recovery probability (γ = 0.4 and 0.04 respectively). Threshold values of each variable are included in each tree. The boxes at the tips of the trees indicate the estimates of average peak time or proportion of the network infected across simulations (top value) and percentage of networks in our dataset to be assigned to this tip. For example, 50% of our empirical networks had spectral radius values ≥ 26 and for these networks we found on average, a maximum of 0.76 of the network infected after 100 time steps. Tip boxes are coloured light to dark blue based on network vulnerability to pathogen spread (e.g., longer time to peak = light blue). Global fit = R^2^ for how well the surrogate model replicates the predictions of the trained model. See Figs. S6 for the complete list of global surrogate models and Fig. S7 for ‘time to peak’ surrogates. Colour of the label indicates what type of predictor it is (blue = spectral, red = non-spectral structural variables, gold = network metadata, see Tables S1 & S2).

**Fig. 5:**
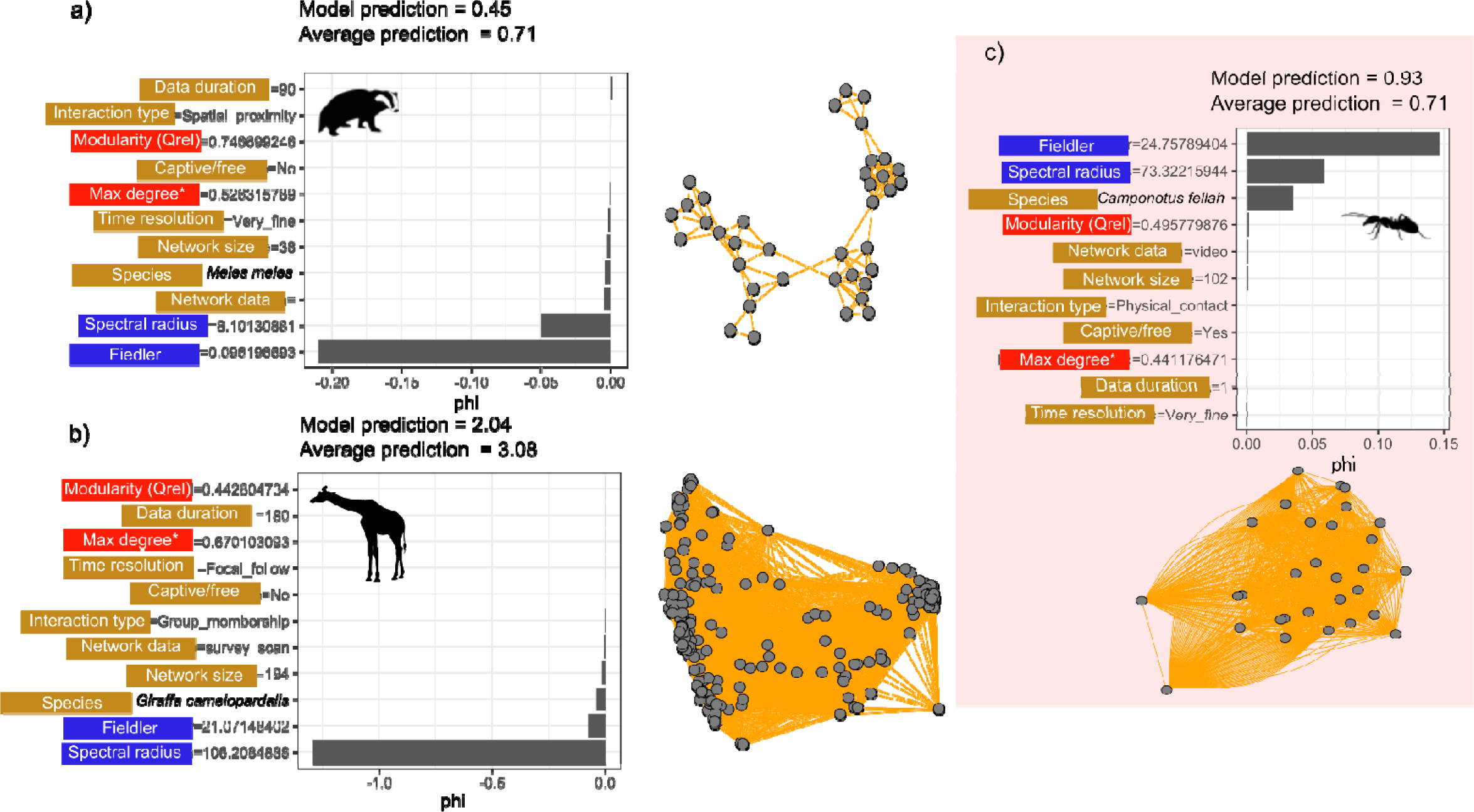
Shapely values (phi) that quantify how each variable shaped a) simulated proportion infected in an empirical badger network, b) simulated time to peak in the empirical giraffe network (VanderWaal *et al*. 2014) and c) simulated proportion infected in an ant network (*C. fel lah*, note nodes overlap) (β = 0.05, γ=0.04). Panels on the right or below are the corresponding networks. Negative Shapely values indicate that the variable reduced time to peak or proportion infection relative to other variables included in the model. Positive Shapely values indicate t that the variables increased the proportion infected relative to other variables included in the model. Colour of the labels indicates what type of predictor it is (blue = spectral, red = non-spectral structural variables, light brown = network metadata, see Tables S2/S3). Corresponding values for each network are next to the label

### Individual networks

To further validate our predictions, we zoomed into how our models predicted spread across individual networks using Shapley values (assuming β = 0.05, γ=0.04). Our models predicted that the European badger network was less vulnerable to disease spread with a much lower prediction of proportion infected compared to the average across all networks (Fig. 5a). This prediction was largely driven by the Fiedler value (0.096, much lower than the mean of 7.31 across all networks) and, to a lesser degree, by the small spectral radius (8.10 compared to a mean of 34.8 across all networks, Fig. 5a). In contrast, our model predicted a faster time to peak for the giraffe network (2.04 timesteps on average compared to the global average of 3.08) and this prediction was driven by the high spectral value (106.02, Fig. 5b) even though the Fiedler value was also high (21.07, Fig. 5b). The Fiedler value was more important in predicting spread across an ant colony which was more vulnerable to disease (Fig. 5c, 93% of the nodes were predicted to be infected on average).

Taken together, our findings show how the spectral values of contact networks offer a valuable and informative “shorthand” for how vulnerable different animal networks are to outbreaks.

## Discussion

Here, we show that the spectral radius and Fiedler value of a network can be a remarkably strong predictor for population vulnerability to diverse epidemics varying in key epidemiological parameters. We demonstrate how a powerful machine learning and simulation approach can effectively predict pathogen outbreak characteristics on a large collection of empirical animal contact networks. We not only demonstrate the high predictive power of a network’s spectral properties but also show that our predictions can help approximate the vulnerability of populations to infectious disease across outbreak scenarios. Further, our findings offer insights into how nuances in social organisation translate into differences in pathogen spread across the animal kingdom.

Across real-world contact networks, we found that the networks’ spectral properties (Fiedler distance and spectral radius) were powerful proxies for pathogen spread. The strong relationship between spectral radius and epidemic threshold has been demonstrated for theoretical networks (Prakash *et al*. 2010) and has been used to assess vulnerability of cattle movement networks to spread of bovine brucellosis (Darbon *et al*. 2018). We expand these findings to show that the spectral radius is the most important predictor in our models of epidemic behaviour across diverse animal social systems. While we examined only SIR propagation through our networks, theoretical results suggest that our findings will extend to other propagation mechanics such as SIS, (susceptible-infected-susceptible) and SEIR (susceptible, exposed, infected, recovered) (Prakash *et al*. 2010).

For some networks and epidemiological parameters, spectral radius alone was not sufficient to predict spread, and the Fiedler value and modularity still played an important role. The Fiedler value and spectral radius of the networks were correlated, but below our ρ = 0.7 threshold (Fig. S2). One potential reason for this is that the Fiedler value seems to be less sensitive to nodes with high connectivity compared to the spectral radius (Fig. 1); however, the mathematical relationship between these two algebraic measures of connectivity is poorly understood (Tang & Priebe 2016). Combined, our global surrogate models and accumulated effects plots pointed to networks with spectral radii > ∼8 and Fiedler values > 1 being more vulnerable to pathogen spread (the effect of the Fiedler value on spread was much weaker overall). The spectral properties were dominant for the fast-spreading pathogen models (e.g., β > 0.1, γ = 0.04), whereas network size and modularity played a more important role in our models for more slowly spreading pathogens (e.g., Figs. S6 & S7).

When modular structure played a role in disease spread in our study, we detected similar patterns to those found by Sah *et al*. (2017). As in Sah *et al*. (2017), we found that epidemic progression was only slowed in highly modular networks (Q_rel_ > ∼0.7) when the probability of transmission between nodes was low (β > 0.025). Such subdivided networks were rare in our data and are commonly associated with high fragmentation (small groups or sub-groups) and high subgroup cohesion (Sah *et al*. 2017). The reduced importance of modularity relative to spectral radius is due to within-group connections being crucial for epidemic outcomes in many contexts (Sah *et al*. 2017). Spectral values may have higher predictive performance, as they summarize connectivity across the networks including between- and within-group connections. Interpreting how modularity alone impacted epidemic outcomes was difficult on these empirical networks, as all modularity measures were strongly correlated with mean degree, diameter and transitivity (Fig. 2, Fig. S2). The extent of these correlations can vary substantially based on other aspects of network structure and they all have interacting effects on disease dynamics (Zhang & Zhang 2009; Ames *et al*. 2011). However, the spectral radius captures epidemiologically important aspects of network structure on its own, without having to untangle whether different aspects of network structure are correlated.

More broadly, our study provides a framework for how interpretable machine learning can predict spread across networks for a wide variety of epidemic parameters. While our RF *MrIML* model had much higher predictive performance compared to the corresponding linear models, further investigation of these models provided critical insight into how network structure impacted pathogen spread. This framework could identify general trends of disease vulnerability, specific thresholds for pathogens with certain characteristics, as well as the drivers of spread for individual networks.

To help practitioners apply our model to different host-pathogen systems, we developed an R-Shiny app (https://spreadpredictr.shinyapps.io/spreadpredictr/). While network data is hard to collect, a wide variety of scientists from behavioural ecologists to livestock and wildlife managers collate network data to address a variety of questions regarding, for example, health and sociality (Craft 2015). Our web application allows users to take a contact network of interest and make predictions of pathogen spread for diverse transmission and recovery probabilities without the need for simulation. While currently limited to pathogens with SIR transmission dynamics, future versions of the application will include, for example, SI and SEIR mechanics. We stress that for practitioners to make accurate predictions for a particular pathogen, contact definitions and the duration of data should be calibrated or multiple thresholds for what constitutes a transmission contact assessed (see Craft 2015). For example, for the giraffe network we included edges that represented individuals seen once together over a period of a year, and predictions of pathogen spread on this network would likely be inflated for pathogens requiring more sustained contact (VanderWaal *et al*. 2014). Nonetheless, this study shows the utility of linking network simulation and interpretable machine learning approaches to tease apart the drivers of spread across empirical wildlife networks.

As this is a broad comparative study of simulated pathogen spread on 603 empirical networks across taxonomic groups, we made important simplifying assumptions. For example, as there were large differences in how the empirical network edges were weighted across taxa (e.g., some networks were weighted by contact duration and others by contact frequency) our approach treated all contacts as equal in unweighted networks, as is done in similar studies (Ames *et al*. 2011; Sah *et al*. 2017). How or if our spectral-based predictions of spread hold on networks with weighted edges remains an open research question. Further, while this study demonstrates the power of repositories such as the ASNR, there are large biases in the taxa covered that must be accounted for in model structure. For example, a large proportion of the networks (∼150) came from one taxon (*C. fellah*). However, removing this one taxon did not qualitatively change our findings. While species identity was important to include in our models to account for the nonindependence of our data, removing species as a predictor in our models also did not alter our findings. Nonetheless, starting to fill in these taxonomic gaps in a systematic way will increase the utility of comparative approaches such as ours and make them generalizable across taxa and populations.

We also simulated spread across static networks, making the assumptions (i) that aggregated networks are representative of social patterns at epidemiologically relevant timescales and (ii) that network change happens more slowly than pathogen spread. While this is reasonable for our study as we are aiming only to identify general rules for how spectral properties influence epidemiological dynamics in animal social networks, it could be problematic when making system-level predictions (Bansal *et al*. 2010). Consequently, including predictions of spread that account for the dynamic nature of contact structure and pathogen-mediated changes in behaviour is an important future extension of this work. However, applying dynamic network models such as temporal exponential random graph models (Krivitsky & Handcock 2014) to estimate spread is computationally demanding and challenging in a comparative setting due to idiosyncrasies in the model-fitting process. While of high predictive value, our models did not capture all aspects of uncertainty. For example, we assumed each network was fully described, with no missing nodes or edges, which is almost always not the case for wildlife studies. How sensitive spectral properties are to missing data is an open question. However, interestingly, removing some edges from ecological networks with high Fiedler values does not appear to strongly impact the stability of the network (Kumar *et al*. 2019).

This paper provides a significant step towards a spectral understanding of pathogen spread in animal networks. We show that the spectral radius of an animal network is a powerful predictor of spread for diverse hosts and pathogens that can be a valuable shortcut for stakeholders to understand the vulnerability of animal networks to disease. We also demonstrate how multivariate interpretable machine learning models can provide novel insights into spread across scales. Moreover, this study has identified the key axes of network structural variation across the animal kingdom that can inform future comparative network research. As rapid advances in location-based tracking and bio-logging (Katzner & Arlettaz 2020) make network data more readily available to wildlife managers, approaches like this one will be of increasing value.

## Authorship statement

NFJ conducted the analysis and wrote the initial draft of the paper to which all authors contributed. RCA helped with some of the coding elements. JR, KVW, MS & RH also provided network data. NFJ, SC, MEC and MC conceived the project.

## Acknowledgements

This project was supported by an Australian Research Council Discovery Project Grant (DP190102020). We would also like to thank Prof. Menna Jones and Prof. Sue VandeWoude for their support and comments on this manuscript.

## Data statement

All data and code to perform the analysis can be found at https://github.com/nfj1380/igraphEpi

