## Supplementary material for "The spectral underpinnings of pathogen spread on animal networks"

**Table S1:** Network structural properties calculated for each network.

| **Structural property** | **Calculation** | **Hypothesized values linked to fast/high spread** | **Type** | **Included in final model?*** |
| --- | --- | --- | --- | --- |
| Spectral radius | Eigenvalue | Large values | Spectral | Y |
| Fiedler value | Eigenvalue | Large values | Spectral | Y |
| Modularity (Q) | Newman’s Q | Small values | Clustering | N |
| Modularity (Qrel) | Q/Qmax | Small values | Clustering | Y |
| Transitivity | Clustering coefficient | Large values | Clustering | N |
| Network size | Number of nodes | Low values | Avg Node characteristic | Y |
| Centralization | Sum of node scores/max node score | Low values | Avg Node characteristic | Y |
| Max degree |  | Large values | Avg Node characteristic | N |
| Max degree (scaled) | Max degree/ network size | Large values | Avg Node characteristic | Y |
| Mean degree | Average number of edges per node | Large values | Avg Node characteristic | N |
| Mean path length | Average number of edges per node | Large values | Avg Node characteristic | N |
| Diameter | Smallest path between most distant nodes | Small values | Avg Node characteristic | N |

*Predictors with correlation coefficients >0.7 were excluded.

**Table S2:** Network metadata recorded for each network.

| **Metadata** | **Details** | **Numeric or Factor** |
| --- | --- | --- |
| Host species/class | Host species and taxonomic class that the network data came from | Factor |
| Interaction type | Do edges reflect direct contact, close spatial proximity, or group membership | Factor |
| Network construction method | Was the data collected using a logger, survey scan or other method**?** | Factor |
| Time resolution | Does the network represent very fine-scale contacts (seconds), fine-scale (minutes), coarse (days/spatial overlap) or focal follow. | Factor |
| Network duration | Over what period was the data collected (days) | Numeric |
| Captive/free? | Was the network recorded for captive or free-ranging populations? | Factor |

**Table S3:** Principal component vector loadings. Largest vector loading for each principal component (PC) are in bold.

|  | **PC1** | **PC2** | **PC3** | **PC4** | **PC5** | **PC6** | **PC7** | **PC8** |
| --- | --- | --- | --- | --- | --- | --- | --- | --- |
| **Fiedler** | 0.26 | 0.19 | 0.18 | -0.17 | **-0.68** | **-0.49** | -0.04 | 0.33 |
| **Spectral_radius** | 0.33 | 0.30 | 0.17 | 0.02 | -0.02 | -0.17 | 0.36 | **-0.60** |
| **Network_size** | 0.11 | 0.52 | 0.26 | -**0.25** | 0.46 | 0.03 | -0.39 | 0.36 |
| **Highest_degree** | 0.17 | **0.57** | -0.25 | -0.02 | 0.11 | 0.04 | 0.00 | -0.28 |
| **highest_degree_scaled** | 0.03 | 0.27 | **-0.79** | 0.20 | -0.22 | 0.11 | -0.08 | 0.19 |
| **Modularity** | -0.38 | 0.16 | 0.07 | -0.07 | -0.19 | 0.06 | -0.08 | -0.23 |
| **Qrel** | -0.32 | 0.19 | -0.04 | 0.12 | 0.28 | -0.33 | **0.71** | 0.38 |
| **Centralization** | 0.12 | 0.03 | -0.59 | 0.59 | -0.22 | 0.40 | -0.23 | 0.03 |
| **Mean_degree** | **0.40** | -0.07 | 0.04 | 0.00 | 0.00 | 0.19 | 0.14 | 0.21 |
| **Transitivity** | 0.34 | -0.01 | 0.08 | -0.15 | -0.11 | 0.62 | 0.40 | 0.19 |
| **Diameter** | -0.35 | 0.24 | 0.16 | -0.14 | -0.25 | 0.32 | 0.11 | -0.01 |
| **Mean_pathLength** | -0.36 | 0.22 | 0.14 | -0.14 | -0.25 | 0.25 | 0.04 | 0.02 |
| **Data_duration_days** | 0.11 | -0.14 | 0.11 | -0.09 | 0.11 | -0.14 | 0.11 | -0.09 |

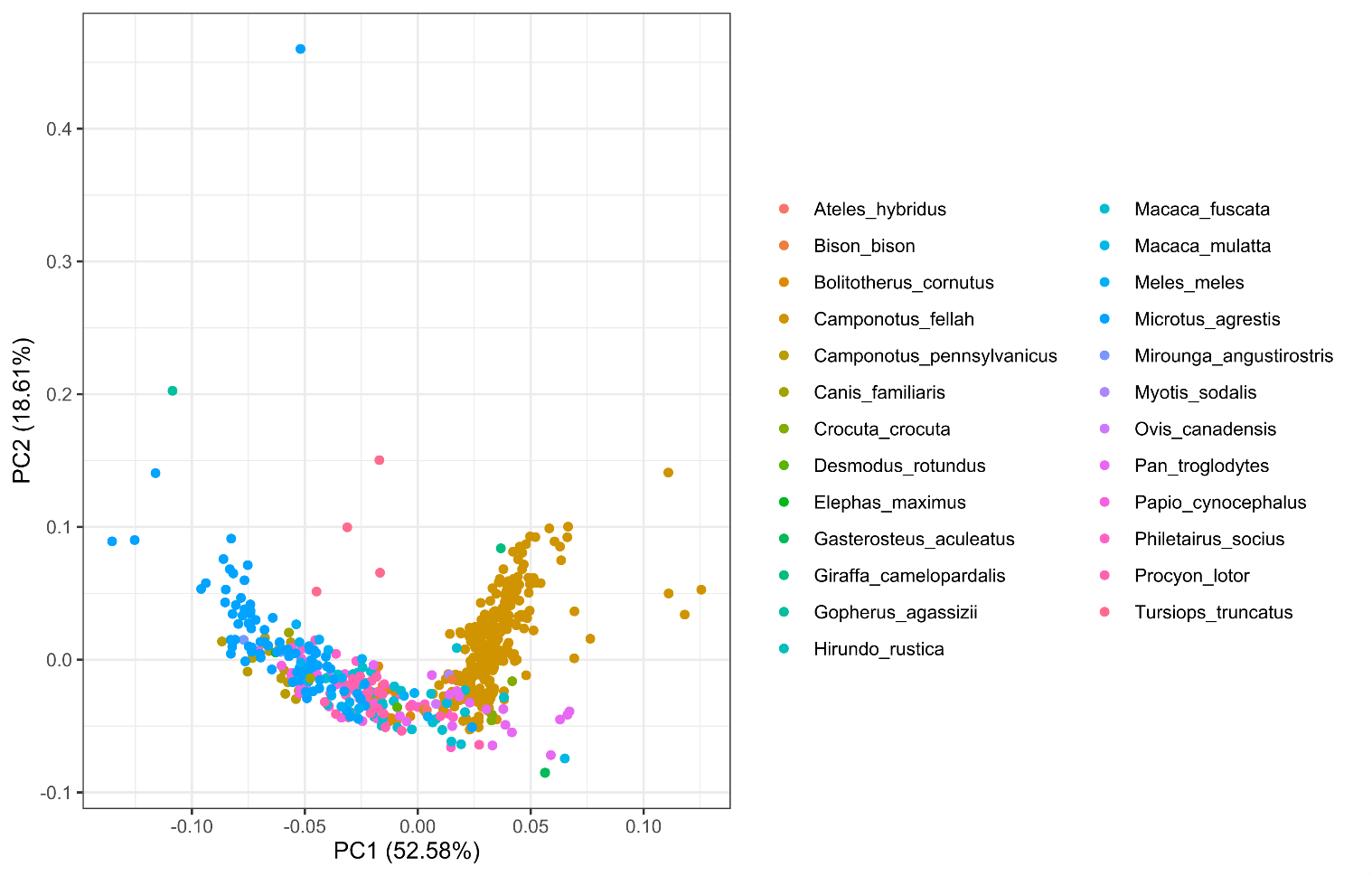

**Fig. S1:** Principal components analysis (PCA) biplot showing that network structure was largely clustered by species. Points are coloured by species with points closer together in Euclidean space being similar in structure and scaled by network size. The PCA was constructed just using continuous network characteristics. Percentages next to PC scores indicate how much variability each axis accounted for in the data.

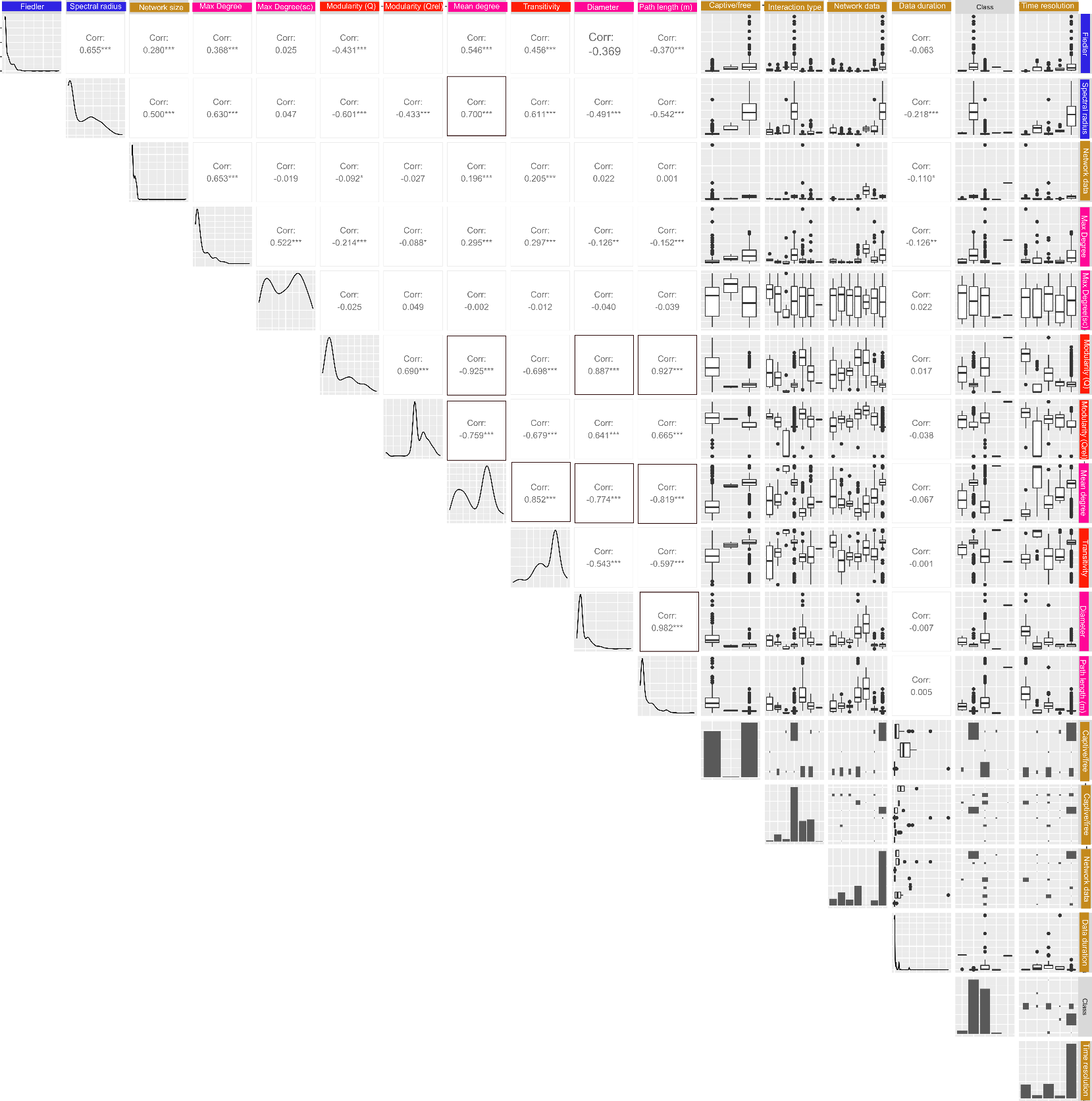

**Fig. S2:** Correlations between network structure predictors (see Table S2). Boxes around correlation coefficients indicate correlations > 0.7 between a pair of predictors. Box and whisker plots are provided for categorical predictors (see Table S3 for details). The diagonal shows the distribution of predictor values

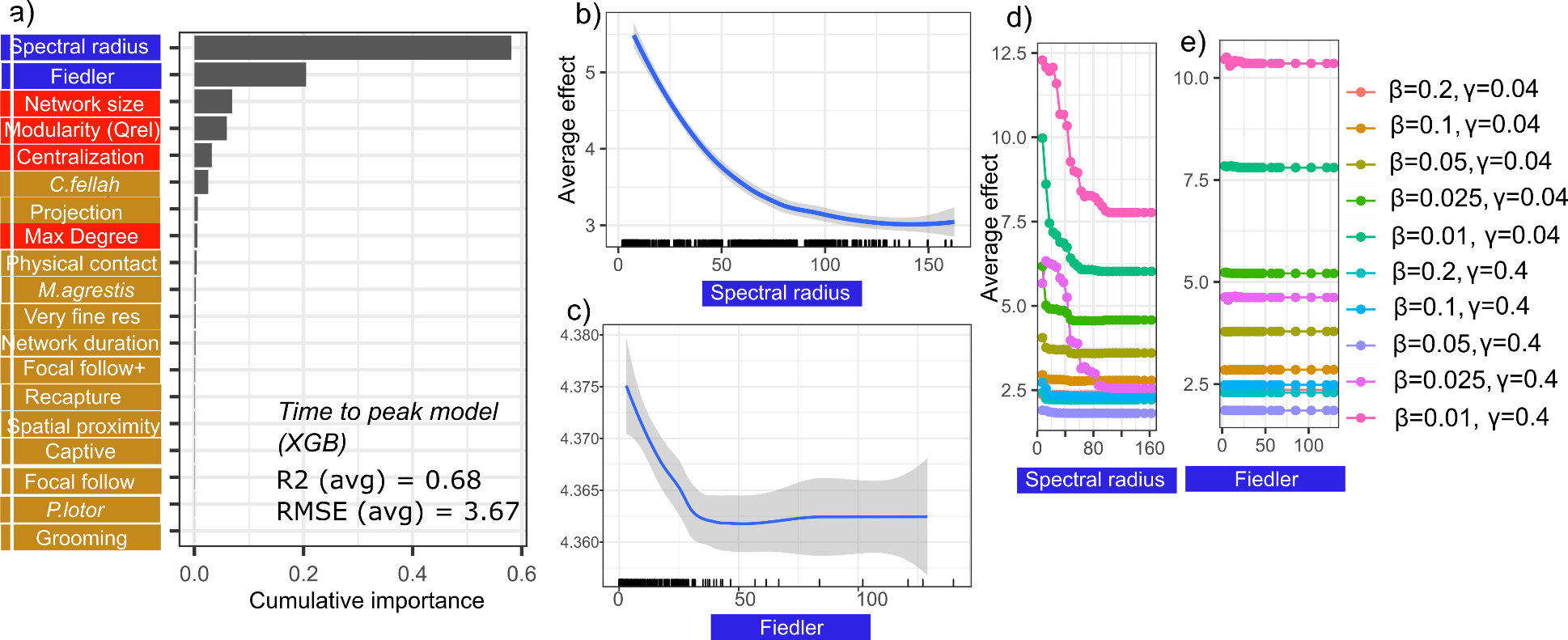

**Fig. S3**: Plots showing the predictive performance, variable importance and the functional form of relationships for our best performing *MrIML* time to peak model (XGB: extreme gradient boosting). a) Spectral radius, Fiedler value, network size and modularity (Qrel) were the most important predictors of time to peak (importance threshold > 0.1). Model performance was less that the proportion infected model (average R^2^ = 0.69 and root mean square error (RMSE) = 3.6, see Table S4 for performance values for all models). Species names are in italics. b-c) Average predictive surface for the top two variables and time to peak across all epidemic values (95% confidence intervals in grey). Rug plot on the x axis shows the distribution of each characteristic across empirical networks. d-e) Accumulated local effects (ALE) plot revealed strongly non-linear relationships between the spectral radius and Fiedler value that varied with SIR model parameters (see the legend). Colour of the labels indicates what type of predictor it is (blue = spectral, red = non-spectral structural variables, gold = network metadata, see Tables S2 & S3). See Fig S3 for the predictive surfaces for modularity and network size.

**Table S4:** Model performance and construction details across algorithms. The model with the highest performance scores (in italics) were further analysed.

| **Model** | **Data** | **R^2^** | **RMSE** | **Hot encoded?**  **(X)** | **Transform (Y)** |
| --- | --- | --- | --- | --- | --- |
| Linear model* | Proportion infected | 0.71 | 0.13 | Yes | NA |
| Support vector machines (SVMs) | Proportion infected | 0.93 | 0.04 | No | NA |
| *Random forests (RF)* | *Proportion infected* | *0.96* | *0.03* | *No* | *NA* |
| RF just using spectral/Fiedler value | Proportion infected | 0.96 | 0.03 | No | NA |
| Extreme gradient boosting  (XGB) | Proportion infected | 0.95 | 0.03 | Yes | NA |
| Linear model | Time to peak | 0.46 | 4.78 | Yes | Log |
| Support vector machines (SVMs) | Time to peak | 0.64 | 3.83 | No | NA |
| Random forests (RF) | Time to peak | 0.68 | 3.52 | No | NA |
| *Extreme gradient boosting*  *(XGB)* | *Time to peak* | *0.69* | *3.67* | *Yes* | *NA* |
| XGB just using spectral/Fiedler value | Time to peak | 0.67 | 3.58 | Yes | NA |

*not optimal for proportional data but currently beta regression is unavailable in the *MrIML* R^2^ and RSME (root mean squared error) estimates are the average of all model combinations

**Table S5:** A comparison of proportion infected estimates across our empirical networks using our machine-learning approach to degree-based mean field approximations (DBMF).

| **β/γ combination** | **RMSE (ML)** | **RMSE (DBMF)** |
| --- | --- | --- |
| 0.2/0.04 (~R_0_=5) | 0.025 | 0.26 |
| 0.1/0.04 (~R_0_=2.5) | 0.03 | 0.33 |
| 0.05/0.04 (~R_0_=1.25) | 0.03 | 0.42 |

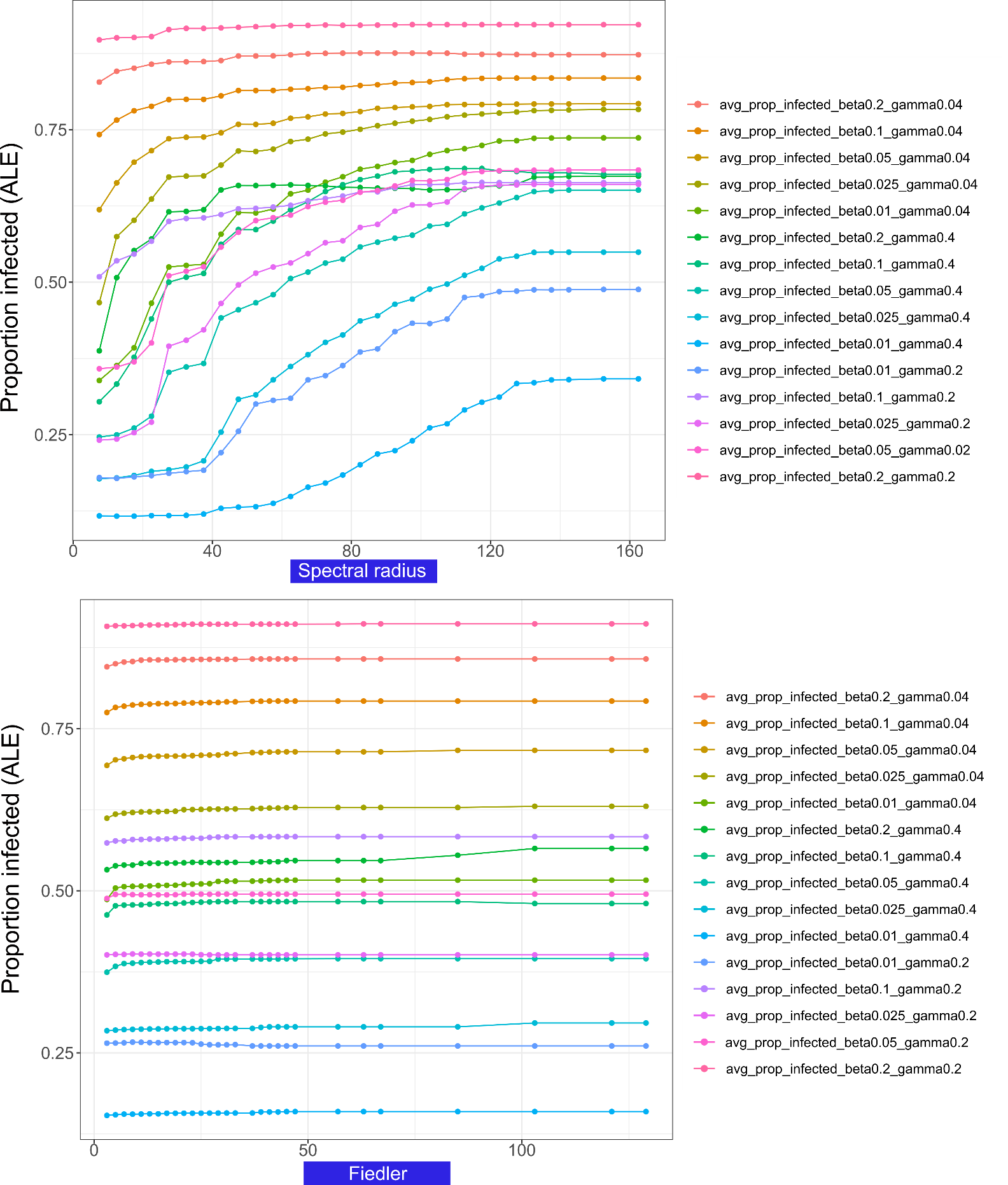

**Fig. S4:** a) Spectral radius and b) Fiedler value accumulated local effects plots including extra recovery rates.

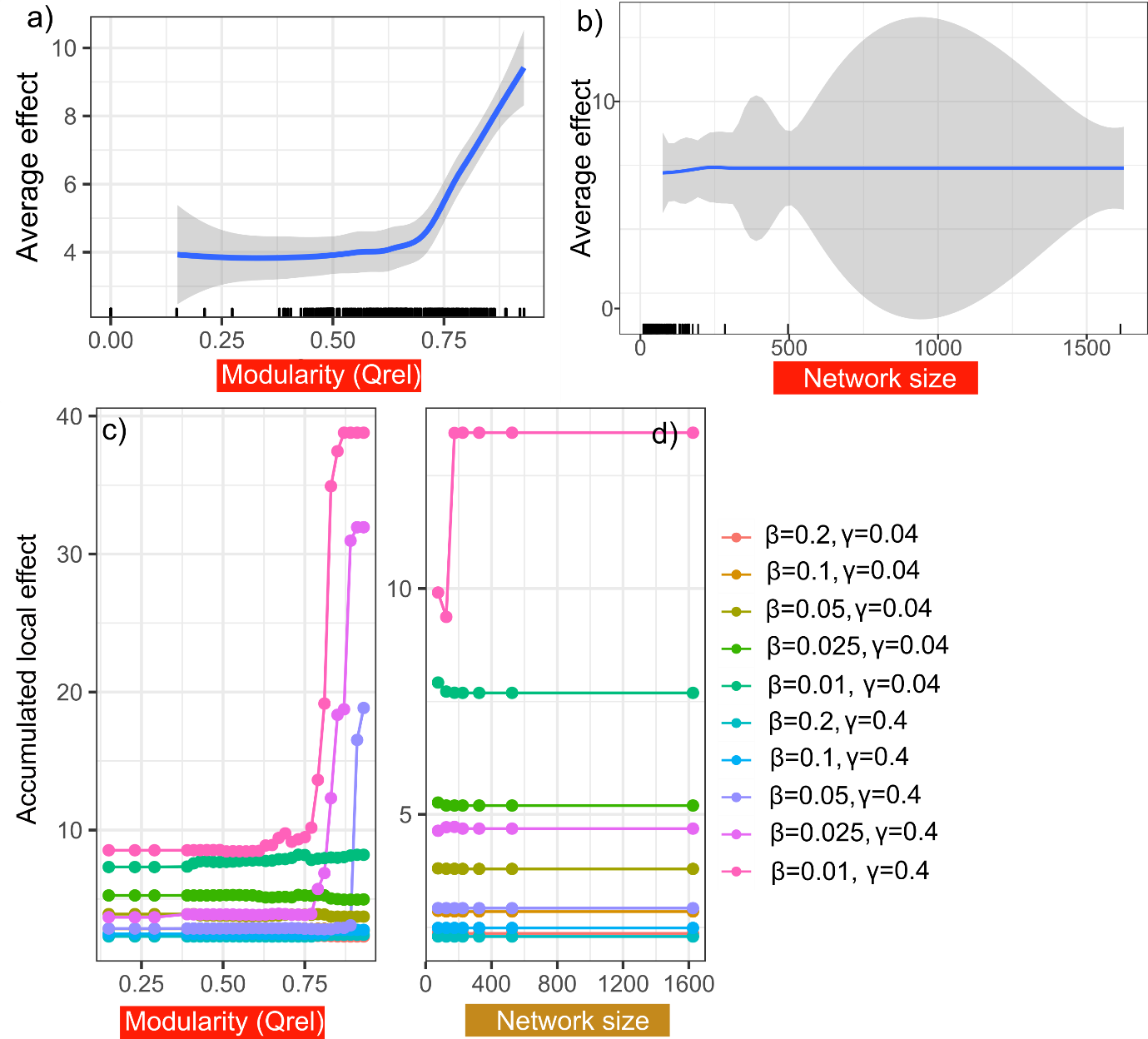

**Fig. S5:** Accumulated local effects (ALE) plots from the extreme gradient boosting model showing the generalized relationship between a) modularity and b) network size on time to epidemic peak. The bottom panels (c & d) show variation across epidemic parameters.

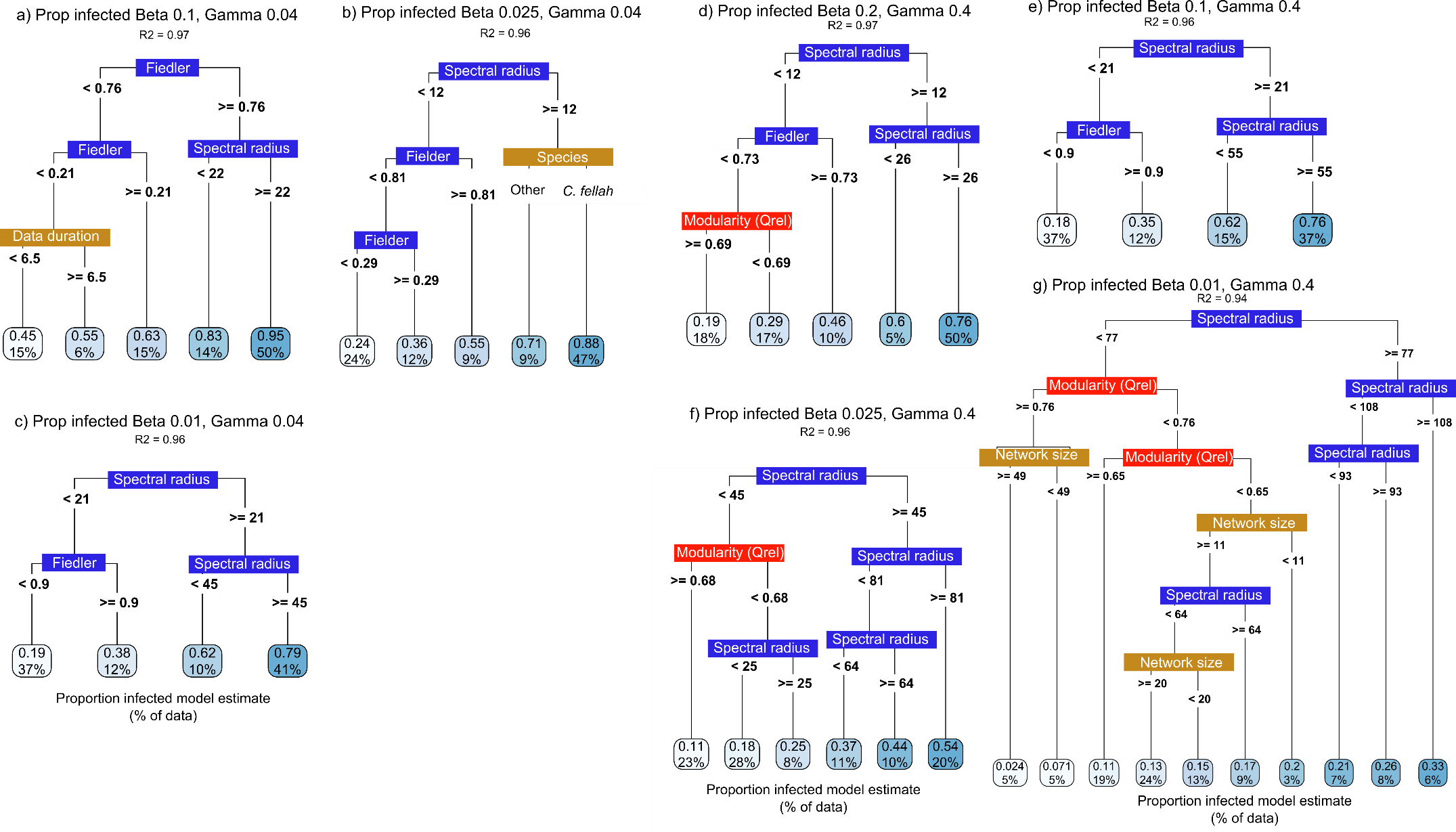

**Fig. S6:** Global surrogate decision trees for all proportion infected models for remaining epidemic parameter sets (see Figure 5 for the other surrogate decision trees). Threshold values of each variable are included in each tree. The boxes at the tips of the trees indicates the estimates proportion of the network infected across simulations (top value) and percentage of networks in our dataset to be assigned to this tip. Tip box are coloured light to dark red based on network vulnerability to pathogen spread (i.e., longer time to peak = light red). Global fit = R^2^ for how well the surrogate model replicates the predictions of the trained model. Colour of the labels indicates what type of predictor it is (blue = spectral, red = clustering, green = network metadata, see Tables S2 & S3).

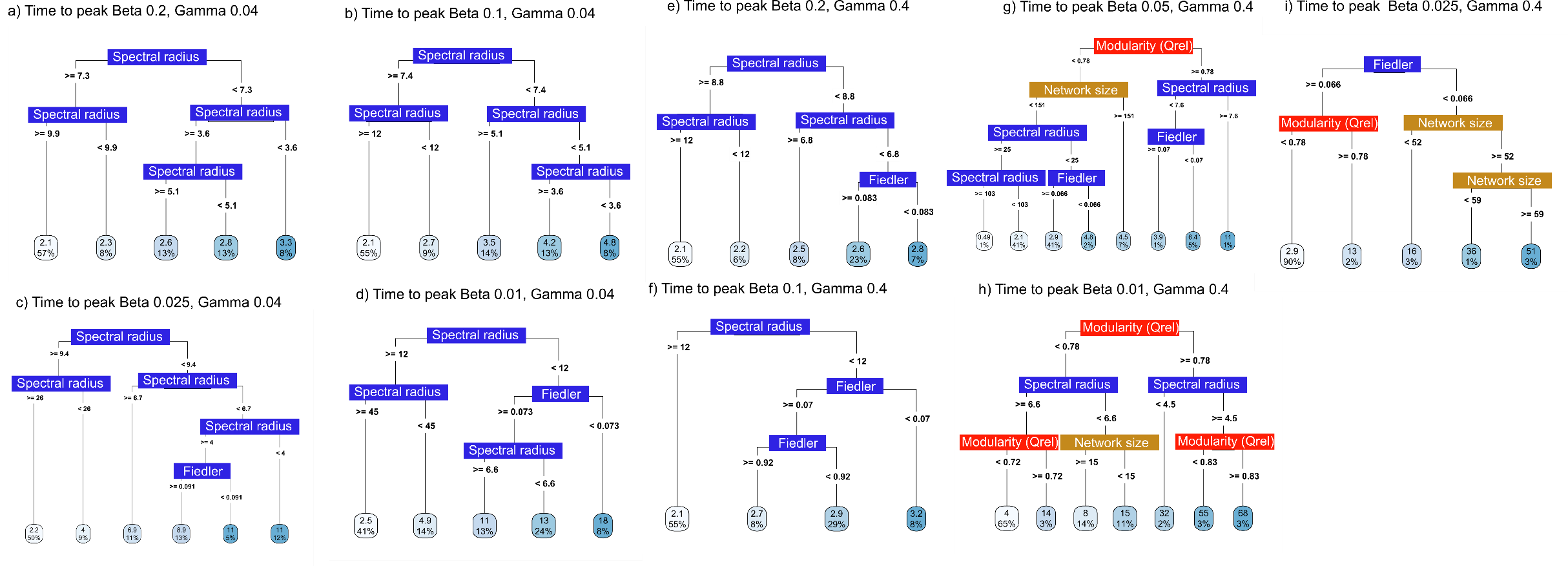

**Fig. S7:** Global surrogate decision trees for all time to peak models for remaining epidemic parameter sets (see Figure for the remaining surrogate decision trees). Threshold values of each variable are included in each tree. The boxes at the tips of the trees indicates the estimates proportion of the network infected across simulations (top value) and percentage of networks in our dataset to be assigned to this tip. Tip box are coloured light to dark red based on network vulnerability to pathogen spread (i.e., longer time to peak = light red). Global fit = R^2^ for how well the surrogate model replicates the predictions of the trained model. Colour of the labels indicates what type of p.
